## Supplemental Information for "Short-term social isolation acts on hypothalamic neurons to promote social behavior in a sex- and context-dependent manner"

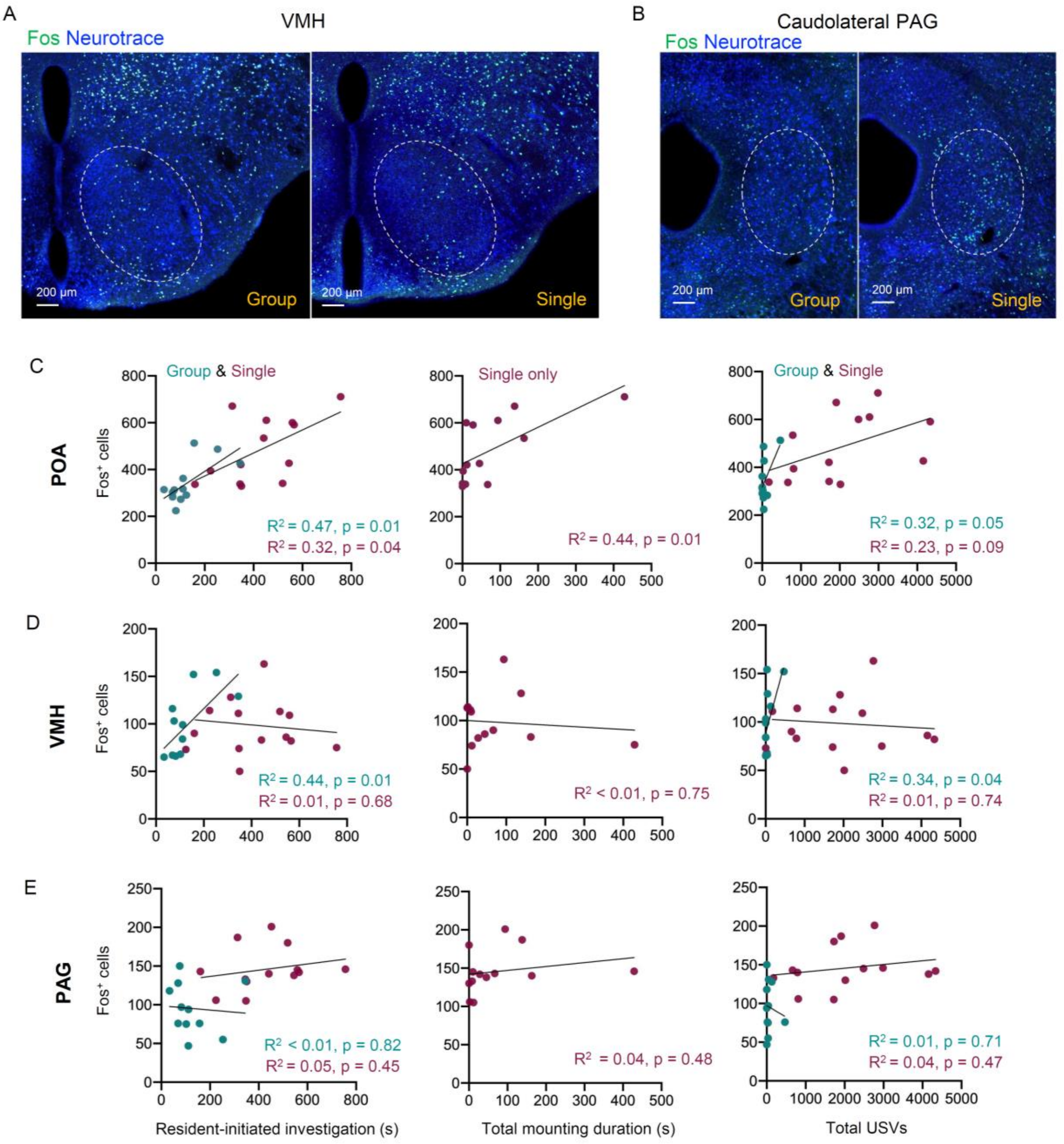


**Figure S1. Additional characterization of Fos expression in single-housed vs. group-housed females and comparison to rates of female social behaviors.** (A) Representative confocal images show Fos expression (green) in the VMH of a group-housed female (left) and a single-housed female (right) following same-sex social interactions. Blue, Neurotrace. (B) Same as (A), for the caudolateral PAG. (C) Left, the relationship between total time spent in resident-initiated social investigation and numbers of Fos-positive POA neurons is shown for group-housed (teal) and single-housed (maroon) female residents following interactions with novel females. Middle, same as left, for total resident-initiated mounting time vs. numbers of Fos-positive POA neurons. Data only shown for single-housed residents because group-housed residents never mounted female visitors. Right, same as left, for total USVs vs. numbers of Fos-positive POA neurons. (D) Left, the relationship between total time spent in resident-initiated social investigation and numbers of Fos-positive VMH neurons is shown for group-housed (teal) and single-housed (maroon) female residents following interactions with novel females. Middle, same as left, for total resident-initiated mounting time vs. numbers of Fos-positive VMH neurons. Right, same as left, for total USVs vs. numbers of Fos-positive VMH neurons. (E) Left, the relationship between total time spent in resident-initiated social investigation and numbers of Fos-positive caudolateral PAG neurons is shown for group-housed (teal) and single-housed (maroon) female residents following interactions with novel females. Middle, same as left, for total resident-initiated mounting time vs. numbers of Fos-positive PAG neurons. Right, same as left, for total USVs vs. numbers of Fos-positive PAG neurons.

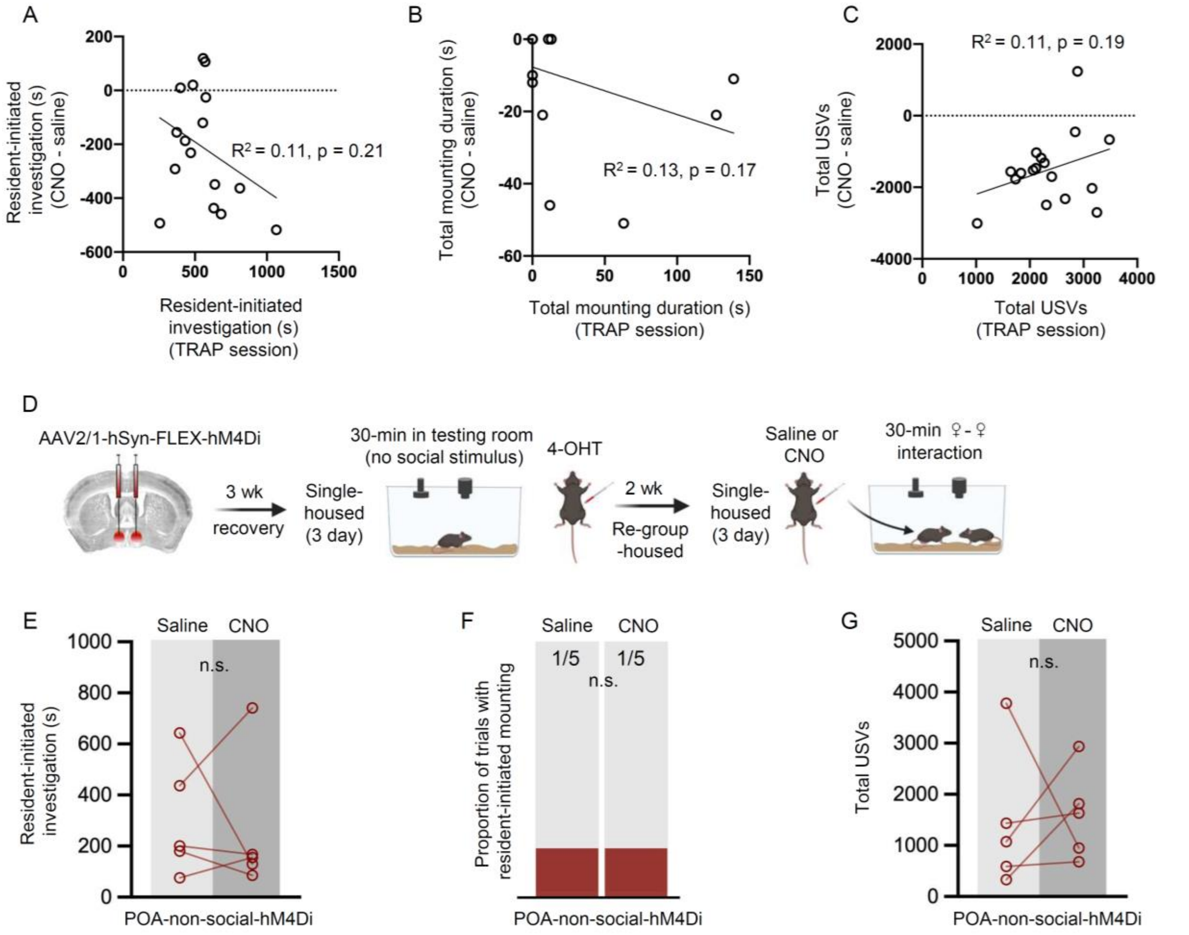


**Figure S2. Additional analyses and control experiments related to Figure 2.** (A) Time spent by POA-social-hM4Di females engaged in social investigation during the TRAPing session is compared to the subsequent change in social investigation in the test sessions (CNO-saline). (B) Same as (A), for time spent mounting. (C) Same as (A), for total USVs. (D) Experimental timeline and viral strategy to chemogenetically inhibit the activity of POA neurons TRAPed in single-housed females that were not given a social interaction. (E) Total time spent in resident-initiated social investigation is shown on saline and CNO days. (F) Same as (E), for proportion of trials with resident-initiated mounting. (G) Same as (E), for total USVs.

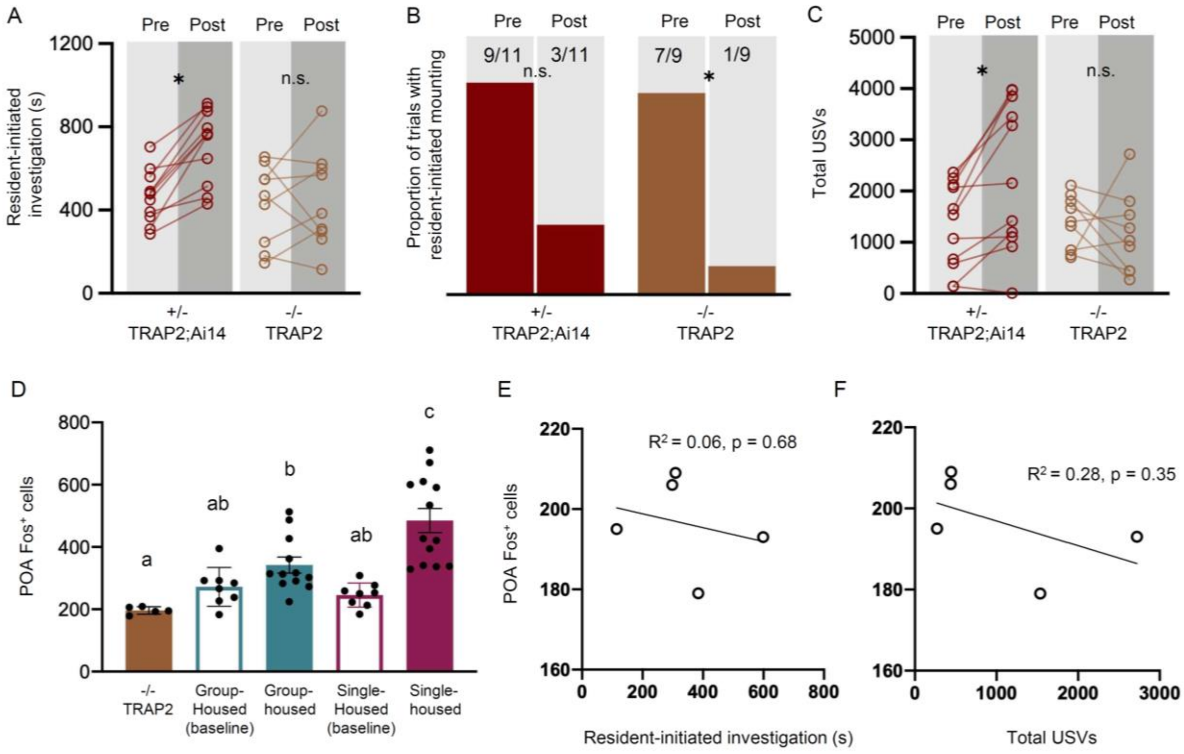


**Figure S3. Comparison of effects of ablation of POA$_{social}$ neurons in TRAP2 heterozygous vs. homozygous females.** (A) Time spent in resident-initiated investigation is show pre- vs. post-4-OHT treatment for TRAP2 heterozygous POA$_{social}$-caspase females (TRAP2;Ai14, red symbols, N = 11) and TRAP2 homozygous POA$_{social}$-caspase females (orange symbols, N = 9). (B) Same as (A), for proportion of trials with resident-initiated mounting. (C) Same as (A), for total USVs. (D) Counts of Fos-positive POA neurons are shown for TRAP2 homozygous POA$_{social}$-caspase females following a same-sex social interaction (orange bar, N = 5), control group-housed females with no social interaction (teal open, N = 8), control group-housed females following a same-sex social interaction (teal filled, N = 12), control single-housed females with no social interaction (maroon open, N = 8), and control single-housed females following a same-sex social interaction (maroon filled, N = 13). Control female groups are the same as those plotted in Fig. 1F. (E) The relationship between total time spent in resident-initiated social investigation and numbers of Fos-positive POA neurons is shown for TRAP2 homozygous POA$_{social}$-caspase females. (F) Same as (E), for the relationship between total USVs and Fos-positive POA neurons.

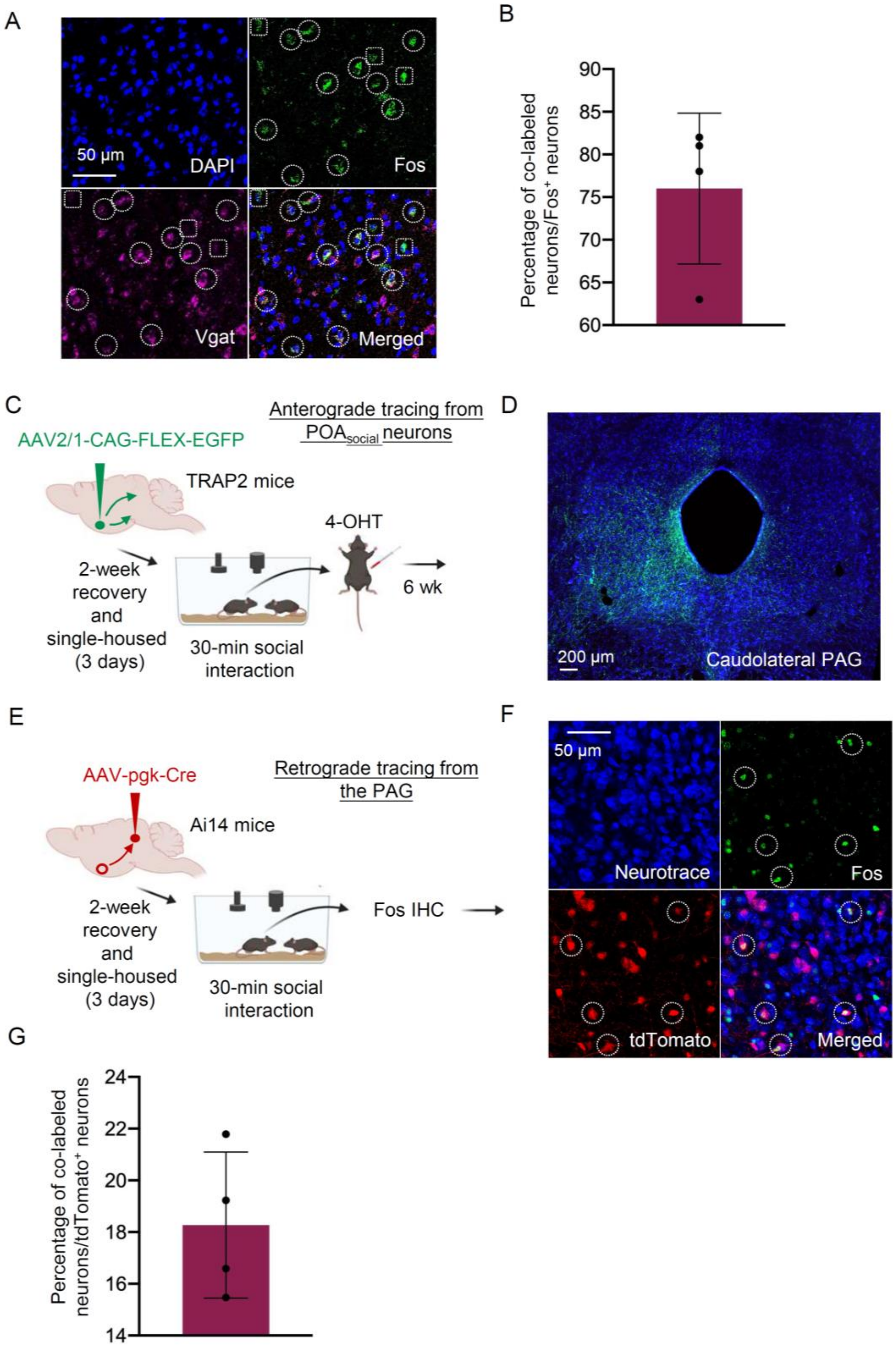


**Figure S4. Characterization of neurotransmitter phenotype and axonal projections of POA$_{social}$ neurons.** (A) Representative confocal images of in situ hybridization performed on brain sections containing the POA, showing overlap of expression of Fos (green) and VGAT (magenta). Blue, DAPI. (B) Quantification of proportion of Fos-positive POA neurons that expressed VGAT. (C) Experimental timeline and viral strategy to express GFP in POA$_{social}$

neurons. (D) Confocal images showing GFP-labeled axons of $POA_{social}$ neurons within the caudolateral PAG. Blue, Neurotrace. (E) Experimental timeline and viral strategy to retrogradely label PAG-projecting POA neurons with tdTomato. (F) Confocal image showing tdTomato labeling in a coronal section containing the POA, and dotted circles in insets indicate examples of double-labeled neurons. Neurotrace, blue. (G) Quantification of proportion of tdTomato-expressing POA neurons that are also Fos-positive.

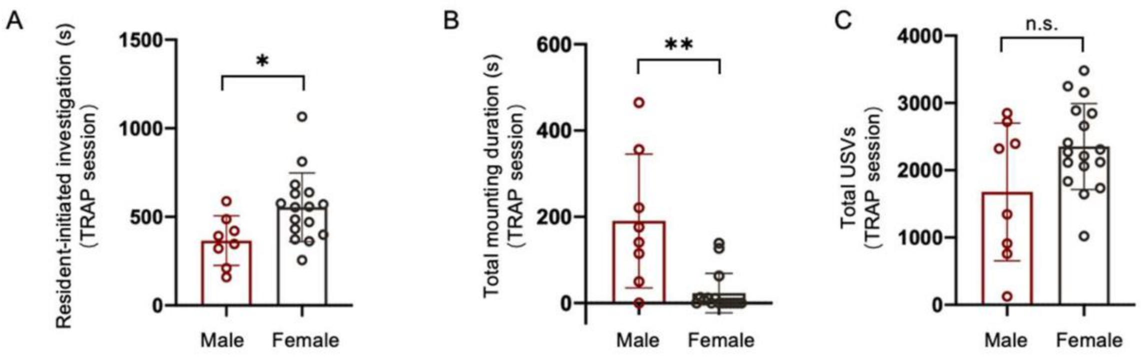


**Figure S5. Comparison of social behavior during TRAPing sessions for male vs. female $POA_{social}$-hM4Di mice.** (A) Total time spent in resident-initiated social investigation during 30-minute TRAPing session interactions with novel females is shown for male (red, N = 9) and female (black, N = 16) $POA_{social}$-hM4Di mice. (B) Same as (A), for time spent in resident-initiated mounting. (C) Same as (A), for total USVs. Please note that N = 2 $POA_{social}$-hM4Di males and N = 1 $POA_{social}$-hM4Di female were excluded because their TRAPing session videos were not saved due to experimenter error.

**Table S1. Details of statistical analyses.**

| **Figure, comparison, and statistical test** | **Group means +/- SD** | **Test results** |
|---|---|---|
| Fig. 1B: total time spent in resident-initiated investigation<br>• T-test | Group-housed residents: 127.8 ± 88.19 (N = 12)<br><br>Single-housed residents: 429.2 ± 161.3 (N =13) | t(23) = -5.72, **P < 0.001** |
| Fig. 1C: proportion of subject females that mounted<br>• Z-test for independent proportions | Group-housed residents = 0 of 12<br><br>Single-housed residents = 11 of 13 | Z = -4.26, **P < 0.001** |
| Fig. 1D: total USVs<br>• Mann-Whitney U test | Pairs with group-housed resident: 64.67 ± 132.0 (N = 12)<br><br>Pairs with single-housed resident: 2042 ± 1292 (N = 13) | Z = 4.16, **P < 0.001** |
| Fig. 1F, left: total number of Fos-positive cells in the POA<br>• Two-way ANOVA (factor 1 = housing; factor 2 = social test); post-hoc Tukey's HSD tests | Group-housed baseline: 272.1 ± 63.3 (N = 8)<br><br>Group-housed social: 342.2 ± 88.7 (N = 12)<br><br>Single-housed baseline: 245.9 ± 38.7 (N = 8)<br><br>Single-housed social: 484.9 ± 139.2 (N = 13) | Main effect of housing: F(1,37) = 3.43, P = 0.07<br>Main effect of social test: F(1,37) = 24.15, **P < 0.01**<br>Interaction F(1,37) = 7.22, **P = 0.01**<br><br>Post-hoc pairwise comparisons:<br>• Group-housed baseline vs. group-housed social: t(37) = -1.56, P = 0.41<br>• Group-housed baseline vs. single-housed baseline: t(37) = 0.54, P = 0.95<br>• Group-housed baseline vs. single-housed social: t(37) = -4.82, **P < 0.001**<br>• Group-housed social vs. single-housed baseline: t(37) = -2.15, P = 0.16<br>• Group-housed social vs. single-housed social: t(37) = -3.63, **P = 0.005**<br>• Single-housed baseline vs. single-housed social: t(37) = -5.42, **P < 0.001** |

| | | |
|---|---|---|
| Fig. 1F, middle: total number of Fos-positive cells in the VMH<br>• Two-way ANOVA (factor 1 = housing; factor 2 = social test) | Group-housed baseline: 86.5 ± 19.8 (N = 8)<br><br>Group-housed social: 98.0 ± 33.2 (N = 12)<br><br>Single-housed baseline: 76.8 ± 17.9 (N = 8)<br><br>Single-housed social: 98.3 ± 28.9 (N = 13) | Main effect of housing: $F(1,37) = 0.29$, $P = 0.59$<br>Main effect of social test: $F(1,37) = 3.63$, $P = 0.06$<br>Interaction $F(1,37) = 0.33$, $P = 0.56$ |
| Fig. 1F, right: total number of Fos-positive cells in the PAG<br>• Two-way ANOVA (factor 1 = housing; factor 2 = social test); post-hoc Tukey's HSD tests | Group-housed baseline: 94.9 ± 19.3 (N = 8)<br><br>Group-housed social: 95.18 ± 33.12 (N = 11)<br><br>Single-housed baseline: 85.1 ± 29.2 (N = 8)<br><br>Single-housed social: 145.8 ± 28.4 (N = 13) | Main effect of housing: $F(1,36) = 4.93$, **$P = 0.03$**<br>Main effect of social test: $F(1,36) = 10.97$, **$P < 0.01$**<br>Interaction $F(1,36) = 10.75$, **$P < 0.01$**<br><br>Post-hoc pairwise comparisons:<br>• Group-housed baseline vs. group-housed social: $t(36) = -0.02$, $P = 1.00$<br>• Group-housed baseline vs. single-housed baseline: $t(36) = 0.95$, $P = 0.90$<br>• Group-housed baseline vs. single-housed social: $t(36) = -3.98$, **$P = 0.002$**<br>• Group-housed social vs. single-housed baseline: $t(36) = -0.76$, $P = 0.87$<br>• Group-housed social vs. single-housed social: $t(36) = -4.34$, **$P < 0.001$**<br>• Single-housed baseline vs. single-housed social: $t(36) = -4.74$, **$P < 0.001$** |
| Fig. 1G, top: effect of re-group-housing on total resident-initiated investigation<br>• Friedman test; post-hoc pairwise Wilcox tests with Bonferroni-corrected p values | Re-grouped, day 0: 103.5 ± 120.0 (N = 6)<br><br>Re-grouped, day 3: 450.0 ± 228.4 (N = 6)<br><br>Re-grouped, day 17: 158.3 ± 67.6 (N = 6) | Main effect of time: X^2(2) = 7, **$P = 0.03$**<br><br>Post-hoc pairwise comparisons:<br>• Day 0 vs. Day 3: $p = 0.03$<br>• Day 0 vs. Day 17: $p = 0.28$<br>• Day 3 vs. Day 17: $p = 0.08$ |

| | | |
|---|---|---|
| Fig. 1G, bottom: effect of 14-days isolation on total resident-initiated investigation<br>• One-way ANOVA with repeated measures; post-hoc Tukey's HSD tests | 14-day-single, day 0:<br>103.5 ± 54.9 (N = 6)<br><br>14-day-single, day 3:<br>543.3 ± 291.9 (N = 6)<br><br>14-day-single, day 17:<br>648.8 ± 241.0 (N = 6) | Main effect of time: F(2,5) = 8.65, **P = 0.02**<br><br>Post-hoc pairwise comparisons:<br>• Day 0 vs. Day 3: t(5) = -3.35, **P = 0.04**<br>• Day 0 vs. Day 17: t(5) = -5.79, **P = 0.005**<br>• Day 3 vs. Day 17: t(5) = 0.59, P = 0.83 |
| Fig. 1H, top: effect of re-group-housing on total resident-initiated mounting time<br>• Friedman test; post-hoc Wilcoxon exact tests with Bonferroni-corrected p values | Re-grouped, day 0:<br>0.17 ± 0.41 (N = 6)<br><br>Re-grouped, day 3:<br>6.67 ± 7.45 (N = 6)<br><br>Re-grouped, day 17:<br>0.83 ± 1.33 (N = 6) | Main effect of time: X^2(2) = 11.99, P = 0.03<br><br>Bonferroni-corrected p value = 0.0167<br>Post-hoc pairwise comparisons:<br>• Day 0 vs. Day 3: W = 3.5, **p = 0.015**<br>• Day 0 vs. Day 17: W = 14, p = 0.45<br>• Day 3 vs. Day 17: W = 30, p = 0.08 |
| Fig. 1H, bottom: effect of 14-days isolation on total resident-initiated mounting<br>• Friedman test | 14-day-single, day 0:<br>0.0 ± 0.0 (N = 6)<br><br>14-day-single, day 3:<br>1.67 ± 2.07 (N = 6)<br><br>14-day-single, day 17:<br>1.5 ± 1.97 (N = 6) | X^2(2) = 5.81, P = 0.32 |
| Fig. 1I, top: effect of re-group-housing on total USVs<br>• One-way ANOVA with repeated measures; post-hoc Tukey's HSD tests | Re-grouped, day 0:<br>179.8 ± 190.2 (N = 6)<br><br>Re-grouped, day 3:<br>2207.2 ± 1486.9 (N = 6)<br><br>Re-grouped, day 17:<br>371.5 ± 235.9 (N = 6) | Main effect of time: F(2,5) = 10.41, **P = 0.02**<br><br>Post-hoc pairwise comparisons:<br>• Day 0 vs. Day 3: t(5) = -3.35, **P = 0.04**<br>• Day 0 vs. Day 17: t(5) = -1.95, P = 0.22<br>• Day 3 vs. Day 17: t(5) = -3.11, P = 0.06 |

| | | |
|---|---|---|
| Fig. 1I, bottom: effect of 14-days isolation on total USVs<br>• One-way ANOVA with repeated measures; post-hoc Tukey's HSD tests | 17-day-single, day 0:<br>177.3 ± 180.2 (N = 6)<br><br>17-day-single, day 3:<br>2089.2 ± 1204.8 (N = 6)<br><br>17-day-single, day 17:<br>2881.2 ± 1694.3 (N = 6) | Main effect of time: $F(2,5) = 16.04$, **P = 0.006**<br><br>Post-hoc pairwise comparisons:<br>• Day 0 vs. Day 3: $t(5) = -4.19$, **P = 0.02**<br>• Day 0 vs. Day 17: $t(5) = -4.16$, **P = 0.02**<br>• Day 3 vs. Day 17: $t(5) = 2.60$, $P = 0.10$ |
| Fig. 1J: Total Fos-positive POA cells<br>• T-test | Single:<br>469.9 ± 45.1 (N = 6)<br><br>Re-grouped:<br>328.4 ± 39.66 (N = 6) | $t(10) = 5.77$, **P < 0.001** |
| Fig. S1C, left: total resident-initiated investigation vs. total Fos-positive POA neurons<br>• Linear regression | See above for group means and standard deviations | Group-housed residents:<br>• $R^2 = 0.47$, $t(10) = 2.98$, **p = 0.01**<br>Single-housed residents:<br>• $R^2 = 0.32$, $t(11) = 2.31$, **p = 0.04** |
| Fig. S1C, middle: total resident-initiated mounting vs. total Fos-positive POA neurons<br>• Linear regression; single-housed only | See above for group means and standard deviations | Single-housed residents:<br>• $R^2 = 0.42$, $t(11) = 2.94$, **p = 0.01** |
| Fig. S1C, right: total USVs vs. total Fos-positive POA neurons<br>• Linear regression | See above for group means and standard deviations | For pairs containing group-housed residents:<br>• $R^2 = 0.32$, $t(10) = 2.21$, $p = 0.05$<br>For pairs containing single-housed residents:<br>• $R^2 = 0.23$, $t(11) = 1.85$, $p = 0.09$ |
| Fig. S1D, left: total resident-initiated investigation vs. total | See above for group means and standard deviations | Group-housed residents:<br>• $R^2 = 0.44$, $t(10) = 2.82$, **p = 0.01**<br>Single-housed residents:<br>• $R^2 = 0.01$, $t(11) = -0.42$, $p = 0.68$ |

| | | |
|---|---|---|
| Fos-positive VMH neurons<br>• Linear regression | | |
| Fig. S1D, middle: total resident-initiated mounting vs. total Fos-positive VMH neurons<br>• Linear regression; single-housed only | See above for group means and standard deviations | Single-housed residents:<br>• $R^2 < 0.01$, $t(11) = -0.32$, $p = 0.75$ |
| Fig. S1D, right: total USVs vs. total Fos-positive VMH neurons<br>• Linear regression | See above for group means and standard deviations | For pairs containing group-housed residents:<br>• $R^2 = 0.34$, $t(10) = 2.28$, **$p = 0.04$**<br>For pairs containing single-housed residents:<br>• $R^2 = 0.01$, $t(11) = -0.34$, $p = 0.74$ |
| Fig. S1E, left: total resident-initiated investigation vs. total Fos-positive PAG neurons<br>• Linear regression | See above for group means and standard deviations | Group-housed residents:<br>• $R^2 = 0.006$, $t(9) = -0.23$, $p = 0.82$<br>Single-housed residents:<br>• $R^2 = 0.05$, $t(11)$ 0.78, $p = 0.45$ |
| Fig. S1E, middle: total resident-initiated mounting vs. total Fos-positive PAG neurons<br>• Linear regression; single-housed only | See above for group means and standard deviations | Single-housed residents:<br>• $R^2 = 0.04$, $t(11) = 0.72$, $p = 0.48$ |
| Fig. S1E, right: total USVs vs. total Fos-positive VMH neurons<br>• Linear regression | See above for group means and standard deviations | For pairs containing group-housed residents:<br>• $R^2 = 0.01$, $t(9) = -0.37$, $p = 0.71$<br>For pairs containing single-housed residents:<br>• $R^2 = 0.04$, $t(11) = 0.74$, $p = 0.47$ |

| | | |
|---|---|---|
| Fig. 2B: effects of chemogenetic inhibition of hypothalamic neurons on resident-initiated investigation<br>• Two-way ANOVA with repeated measures on one factor (between-subjects factor = group, within-subjects factor = drug); post-hoc Tukey's HSD tests | POA-social-hM4Di, saline: 661.0 ± 233.5 (N = 17)<br><br>POA-social-hM4Di, CNO: 437.10 ± 242.9 (N =17)<br><br>POA-social-GFP, saline: 581.5 ± 201.4 (N =14)<br><br>POA-social-GFP, CNO: 560.4 ± 180 (N =14)<br><br>AH-TRAPed-hM4Di, saline:<br>474.1 ± 122.5 (N = 12)<br><br>AH-TRAPed-hM4Di, CNO:<br>488.9 ± 120.3 (N =12)<br><br>VMH-TRAPed-hM4Di, saline: 430.8 ± 137.5 (N =5)<br><br>VMH-TRAPed-hM4Di, CNO: 613.5 ± 115.1 (N =5) | Main effect of group: F(3,44) = 0.85, P = 0.47<br>Main effect of drug: F(1,44) = 0.05, P = 0.83<br>Interaction: F(3,44) = 6.73, **P < 0.001**<br><br>Within group post-hoc pairwise comparisons:<br>• POA-social-hM4Di, saline vs. CNO: t(44) = -4.43, **P = 0.0015**<br>• POA-social-GFP, saline vs. CNO: t(44) = -0.38, P = 1.00<br>• AH-TRAPed-hM4Di, saline vs. CNO: t(44) = 0.56, P = 1.00<br>• VMH-TRAPed-hM4Di, saline vs. CNO: t(44) = 1.96, P = 0.52<br><br>Across group post-hoc pairwise comparisons (saline vs. saline or CNO vs. CNO only):<br>• AH-TRAPed-hM4Di CNO vs. POA-social-GFP CNO: t(44) = -0.93, P = 0.98<br>• AH-TRAPed-hM4Di CNO vs. POA-social-hM4Di CNO: t(44) = 0.76, P = 0.99<br>• AH-TRAPed-hM4Di CNO vs. VMH-TRAPed-hM4Di CNO: t(44) = -1.21, P = 0.92<br>• POA-social-GFP CNO vs. POA-social-hM4Di CNO: t(44) = 1.81, P = 0.62<br>• POA-social-GFP CNO vs. VMH-TRAPed-hM4Di CNO: t(44) = -0.54, P = 1.00<br>• POA-social-hM4Di CNO vs. VMH-TRAPed-hM4Di CNO: t(44) = -1.84, P = 0.60<br>• AH-TRAPed-hM4Di saline vs. POA-social-GFP saline: t(44) = -1.62, P = 0.74<br>• AH-TRAPed-hM4Di saline vs. POA-social-hM4Di saline: t(44) = -2.77, P = 0.12<br>• AH-TRAPed-hM4Di saline vs. VMH-TRAPed-hM4Di saline: t(44) = 0.26, P = 1.00<br>• POA-social-GFP saline vs. POA-social-hM4Di saline: t(44) = -1.13, P = 0.95<br>• POA-social-GFP saline vs. VMH-TRAPed-hM4Di saline: t(44) = 1.49, P = 0.81<br>• POA-social-hM4Di saline vs. VMH-TRAPed-hM4Di saline: t(44) = 2.33, P = 0.30 |
| Fig. 2C: effects of chemogenetic inhibition of hypothalamic neurons | POA-social-hM4Di:<br>7 of 17 on saline day,<br>1 of 17 on CNO day | POA-social-hM4Di, CNO vs. saline: X^2 (1) = 4.17, p = **0.04** |

| | | |
|---|---|---|
| on proportion of trials with resident-initiated mounting<br>• McNemar's test for paired proportions | POA-social-GFP:<br>7 of 14 on saline day,<br>7 of 14 on CNO day<br><br>AH-TRAPed-hM4Di:<br>6 of 12 on saline day,<br>3 of 12 on CNO day<br><br>VMH-TRAPed-hM4Di:<br>4 of 5 on saline day,<br>4 of 5 on CNO day | POA-social-GFP, CNO vs. saline: X^2 (1) = 0.25, p = 0.62<br><br>AH-TRAPed-hM4Di, CNO vs. saline: X^2 (1) = 0.57, p = 0.45<br><br>VMH-TRAPed-hM4Di, CNO vs. saline: X^2 (1) = 0, p = 1.00 |
| Fig. 2D: effects of chemogenetic inhibition of hypothalamic neurons on total USVs<br>• Two-way ANOVA with repeated measures on one factor (between-subjects factor = group, within-subjects factor = drug); post-hoc Tukey's HSD tests | POA-social-hM4Di, saline: 2756 ± 802.7 (N = 17)<br><br>POA-social-hM4Di, CNO: 1250 ± 1080.9 (N =17)<br><br>POA-social-GFP, saline: 2221 ± 1034 (N =14)<br><br>POA-social-GFP, CNO: 2225 ± 1110 (N =14)<br><br>AH-TRAPed-hM4Di, saline:<br>1694 ± 638.3 (N = 12)<br><br>AH-TRAPed-hM4Di, CNO:<br>1707 ± 783.5 (N =12)<br><br>VMH-TRAPed-hM4Di, saline: 1640 ± 771.8 (N =5)<br><br>VMH-TRAPed-hM4Di, CNO: 2180 ± 599.6 (N =5) | Main effect of group: F(3,44) = 0.95, P = 0.42<br>Main effect of drug: F(1,44) = 2.58, P = 0.11<br>Interaction: F(3,44) = 11.59, **P < 0.001**<br><br>Within group post-hoc pairwise comparisons:<br>• POA-social-hM4Di, saline vs. CNO: t(44) = -6.77, **P < 0.001**<br>• POA-social-GFP, saline vs. CNO: t(44) = 0.017, P = 1.00<br>• AH-TRAPed-hM4Di, saline vs. CNO: t(44) = 0.048, P = 1.00<br>• VMH-TRAPed-hM4Di, saline vs. CNO: t(44) = 1.32, P = 0.89<br><br>Across group post-hoc pairwise comparisons (saline vs. saline or CNO vs. CNO only):<br>• AH-TRAPed-hM4Di CNO vs. POA-social-GFP CNO: t(44) = -1.33, P = 0.88<br>• AH-TRAPed-hM4Di CNO vs. POA-social-hM4Di CNO: t(44) = 1.23, P = 0.92<br>• AH-TRAPed-hM4Di CNO vs. VMH-TRAPed-hM4Di CNO: t(44) = -0.90, P = 0.98<br>• POA-social-GFP CNO vs. POA-social-hM4Di CNO: t(44) = 2.74, P = 0.14<br>• POA-social-GFP CNO vs. VMH-TRAPed-hM4Di CNO: t(44) = 0.09, P = 1.00<br>• POA-social-hM4Di CNO vs. VMH-TRAPed-hM4Di CNO: t(44) = -1.85, P = 0.59<br>• AH-TRAPed-hM4Di saline vs. POA-social-GFP saline: t(44) = -1.59, P = 0.75 |

| | | |
|---|---|---|
| | | • AH-TRAPed-hM4Di saline vs. POA-social-hM4Di saline: t(44) = -3.35, **P = 0.03**<br>• AH-TRAPed-hM4Di saline vs. VMH-TRAPed-hM4Di saline: t(44) = -0.12, P = 1.00<br>• POA-social-GFP saline vs. POA-social-hM4Di saline: t(44) = -1.77, P = 0.65<br>• POA-social-GFP saline vs. VMH-TRAPed-hM4Di saline: t(44) = 1.33, P = 0.89<br>• POA-social-hM4Di saline vs. VMH-TRAPed-hM4Di saline: t(44) = 2.61, P = 0.18 |
| Fig. 2E: total movement of POA-social-hM4Di females<br>• Paired t-test | POA-social-hM4DI, saline:<br>12509 ± 2550 (N = 17)<br><br>POA-social-hM4Di, CNO:<br>13257 ± 3503 (N = 17) | t(16) = -1.01, P = 0.33 |
| Fig. S2A: total TRAPing session resident-initiated investigation vs. change in resident-initiated investigation (CNO-saline) in test sessions<br>• Linear regression | See above for group means and standard deviations | $R^2$ = 0.11, t(14) = -1.32, p = 0.21 |
| Fig. S2B: total TRAPing session resident-initiated mounting vs. change in resident-initiated mounting (CNO-saline) in test sessions<br>• Linear regression | See above for group means and standard deviations | $R^2$ = 0.13, t(14) = -1.45, p = 0.17 |
| Fig. S2C: total TRAPing session USVs vs. change in USVs (CNO-saline) in test sessions<br>• Linear regression | See above for group means and standard deviations | $R^2$ = 0.11, t(15) = -1.45, p = 0.19 |

| | | |
|---|---|---|
| Fig. S2E: effects of chemogenetic inhibition of POA neurons on resident-initiated investigation, non-social control females<br>• Paired t-test | POA-non-social-hM4Di, saline:<br>307.4 ± 229.4 (N = 5)<br><br>POA-non-social-hM4Di, CNO:<br>256.0 ± 272.9 (N =5) | T(4) = 0.38, p = 0.72 |
| Fig. S2F: effects of chemogenetic inhibition of POA neurons on proportion of trials with resident-initiated mounting, non-social control females<br>• McNemar's test for paired proportions | Saline: 1 of 5<br><br>CNO: 1 of 5 | CNO vs. saline: X^2 (1) = 0, p = 1.00 |
| Fig. S2G: effects of chemogenetic inhibition of POA neurons on total USVs, non-social control females<br>• Paired t-test | POA-non-social-hM4Di, saline:<br>1442 ± 1375 (N = 5)<br><br>POA-non-social-hM4Di, CNO:<br>1603 ± 879.0 (N = 5) | T(4) = -0.2, p = 0.85 |
| Fig. 3B: effects of caspase-mediated ablation of POA neurons on resident-initiated investigation<br>• Two-way ANOVA with repeated measures on one factor (between-subjects factor = group, within-subjects factor = time) | POA-social-caspase, pre-4-OHT:<br>489.0 ± 119.0 (N = 15)<br><br>POA-social-caspase, post-4-OHT: 688.8 ± 215.1 (N = 15)<br><br>POA-social-GFP, pre-4-OHT:<br>476.3 ± 155.9 (N = 13)<br><br>POA-social-GFP, post-4-OHT: 545.4 ± 155.4 (N = 13) | Main effect of group: F(1,26) = 2.84, P = 0.10<br>Main effect of time: F(1,26) =10.08, **P = 0.004**<br>Interaction: F(1,26) = 2.38, P = 0.14 |

| | | |
|---|---|---|
| Fig. 3C: effects of caspase-mediated ablation of POA neurons on proportion of trials with resident-initiated mounting<br>• McNemar's test for paired proportions | POA-social-caspase:<br>13 of 15 on saline day,<br>4 of 15 on CNO day<br><br>POA-social-GFP:<br>8 of 13 on saline day,<br>6 of 13 on CNO day | POA-social-caspase, CNO vs. saline: X^2 (1) = 4.92, p = **0.03**<br><br>POA-social-GFP, CNO vs. saline: X^2 (1) = 0.50, p = 0.48 |
| Fig. 3D: effects of caspase-mediated ablation of POA neurons on total USVs<br>• Two-way ANOVA with repeated measures on one factor (between-subjects factor = group, within-subjects factor = time); post-hoc Tukey's HSD tests | POA-social-caspase, pre-4-OHT:<br>1411.8 ± 771.6 (N = 15)<br><br>POA-social-caspase, post-4-OHT:<br>2024.3 ± 1325.1 (N =15)<br><br>POA-social-GFP, pre-4-OHT:<br>2562.9 ± 1063.9 (N = 13)<br><br>POA-social-GFP, post-4-OHT:<br>2152.9 ± 1043.3 (N =13) | Main effect of group: F(1,26) = 3.11, P = 0.09<br>Main effect of time: F(1,26) =0.31, P = 0.58<br>Interaction: F(1,26) = 7.98, **P < 0.01**<br><br>Post-hoc pairwise comparisons:<br>POA-social-caspase pre vs. POA-social-caspase-post: t(26) = 2.48, P = 0.09<br>POA-social-caspase pre vs. POA-social-GFP-pre: t(26) = -3.31, **P = 0.01**<br>POA-social-caspase pre vs. POA-social-GFP-post: t(26) = 1.81, P = 0.29<br>POA-social-caspase post vs. POA-social-GFP-pre: t(26) = -1.34, P = 0.55<br>POA-social-caspase post vs. POA-social-GFP-post: t(26) = -0.28, P = 0.99<br>POA-social-GFP pre vs. POA-social-GFP-post: t(26) = -1.55, P = 0.43 |
| Fig. S3A: effects of caspase-mediated ablation of POA neurons on resident-initiated investigation<br>• Two-way ANOVA with repeated measures on one factor (between-subjects factor = group, within-subjects factor = time) | TRAP2 heterozygous, pre-4-OHT:<br>465.0 ± 125.3 (N = 11)<br><br>TRAP2 heterozygous, post-4-OHT:<br>711.1 ± 173.8 (N =11)<br><br>TRAP2 homozygous, pre-4-OHT:<br>428.2 ± 195.5 (N = 9)<br><br>TRAP2 homozygous, post-4-OHT: | Main effect of group: F(1,18) = 4.26, P = 0.054<br>Main effect of time: F(1,18) =12.06, **P = 0.003**<br>Interaction: F(1,18) = 8.76, **P < 0.01**<br><br>Within group post-hoc pairwise comparisons:<br>Heterozygous pre vs. heterozygous post: t(18) = 4.80, **P < 0.001**<br>Homozygous pre vs. homozygous post: t(18) = 0.35, P = 0.99<br><br>Across group post-hoc pairwise comparisons:<br>Heterozygous post vs. homozygous post: t(18) = 2.88, **P = 0.045**<br>Heterozygous post vs. homozygous pre: t(18) = 3.49, **P = 0.01** |

| | | |
|---|---|---|
| | 447.8 ± 235.5 (N =9) | Homozygous post vs. heterozygous pre: t(18) = -0.21, P = 1.00<br>Heterozygous pre vs. Homozygous pre: t(18) = 0.51, P = 0.95 |
| Fig. S3B: effects of caspase-mediated ablation of POA neurons on proportion of trials with resident-initiated mounting, non-social control females<br>• McNemar's test for paired proportions | TRAP2 heterozygous:<br>9 of 11 on saline day,<br>3 of 11 on CNO day<br><br>TRAP2 homozygous:<br>7 of 9 on saline day,<br>1 of 9 on CNO day | TRAP2 heterozygous, pre-4-OHT vs. post-4-OHT: X^2 (1) = 2.5, p = 0.11<br><br>TRAP2 homozygous, pre-4-OHT vs. post-4-OHT: X^2 (1) = 4.17, **p = 0.04** |
| Fig. S3C: effects of caspase-mediated ablation of POA neurons on total USV<br>• Two-way ANOVA with repeated measures on one factor (between-subjects factor = group, within-subjects factor = time) | TRAP2 heterozygous, pre-4-OHT:<br>1330 ± 844.0 (N = 11)<br><br>TRAP2 heterozygous, post-4-OHT:<br>2303 ± 1446 (N =11)<br><br>TRAP2 homozygous, pre-4-OHT:<br>1392 ± 526.1 (N = 9)<br><br>TRAP2 homozygous, post-4-OHT:<br>1160 ± 782.2 (N =9) | Main effect of group: F(1,18) = 1.88, P = 0.19<br>Main effect of time: F(1,18) = 3.28, P = 0.09<br>Interaction: F(1,18) = 8.70, **P < 0.01**<br><br>Within group post-hoc pairwise comparisons:<br>Heterozygous pre vs. heterozygous post: t(18) = 3.55, **P = 0.01**<br>Homozygous pre vs. homozygous post: t(18) = -0.77, P = 0.87<br><br>Across group post-hoc pairwise comparisons:<br>Heterozygous post vs. homozygous post: t(18) = 2.12, P = 0.18<br>Heterozygous post vs. homozygous pre: t(18) = 2.10, P = 0.19<br>Homozygous post vs. heterozygous pre: t(18) = -0.21, P = 1.00<br>Heterozygous pre vs. Homozygous pre: t(18) = 0.51, P = 0.95 |
| Fig. S3D: comparison of counts of Fos-positive POA neurons in female groups following same-sex interactions:<br>• One-way ANOVA; post- | TRAP2 -/- POA-social caspase:<br>196.4 ± 11.9 (N =5)<br><br>Group-housed baseline:<br>272.1 ± 63.3 (N = 8) | Main effect of group: F(4,41) = 13.88, **P < 0.001**<br><br>Post-hoc comparisons:<br>Caspase vs. GH baseline: t(41) = -1.42, P = 0.62<br>Caspase vs. GH social: t(41) = 2.93, **P = 0.04**<br>Caspase vs. SH social: t(41) = -0.93, P = 0.88<br>Caspase vs. SH social: t(41) = -5.87, **P < 0.0001** |

| | | |
|---|---|---|
| hoc Tukey's HSD tests | Group-housed social: 342.2 ± 88.7 (N = 12)<br><br>Single-housed baseline: 245.9 ± 38.7 (N = 8)<br><br>Single-housed social: 484.9 ± 139.2 (N = 13) | GH baseline vs. GH social: t(41) = -1.64, P = 0.48<br>GH baseline vs. SH baseline: t(41) = -0.56, P = 0.98<br>GH baseline vs. SH social: t(41) = -5.07, **P = 0.0001**<br>GH social vs. SH baseline: t(41) = 2.26, P = 0.18<br>GH social vs. SH social: t(41) = -3.82, **P = 0.004**<br>SH baseline vs. SH social: t(41) = -5.70, **P < 0.0001** |
| Fig. S3E: total resident-initiated investigation vs. total Fos-positive POA neurons | See above for group means and standard deviations | R^2 = 0.06, t(3) = -0.46, p = 0.68 |
| Fig. S3F: total USVs vs. total Fos-positive POA neurons | See above for group means and standard deviations | R^2 = 0.28, t(3) = -1.09, p = 0.35 |
| Fig. 4B: USVs per second, solo sessions<br>• Mann Whitney U test performed on the difference in USV rates | POA-social-ChR2, laser – pre-laser:<br>1.43 ± 2.49 (N = 9)<br><br>POA-social-GFP, laser - pre-laser: 0.0 ± 0.0 (N =6) | Z = 1.71, P = 0.09 |
| Fig. 4C: USVs per second, social sessions<br>• Mann Whitney U test performed on the difference in USV rates | POA-social-ChR2, laser - pre-laser:<br>1.98 ± 1.74 (N = 9)<br><br>POA-social-GFP, laser - pre-laser:<br>0.04 ± 0.09 (N = 6) | Z = 2.77, **P = 0.006** |
| In text, related to Fig. 4C: USVs per second, laser vs. pre-laser, according to distance between females at time of optogenetic activation<br>• Paired t-test | "Near" stimulations for POA-iso-ChR2 mice, laser – pre-laser:<br>2.97 ± 1.32 (N = 7)<br><br>"Far" stimulations for POA-iso-ChR2 mice, laser – pre-laser: 1.84 ± 1.75 (N = 7) | t(6) = 3.07, **P = 0.02** |

| | | |
|---|---|---|
| | N = 2 females excluded that did not have “far” stimulations | |
| Fig. 4E: percentage of laser stimulations followed by social investigation<br>• T-test | POA-social-ChR2:<br>41.5 ± 31.2 (N = 9)<br><br>POA-social-GFP:<br>10.8 ± 5.0 (N = 6) | T(13) = 2.36, **p = 0.03** |
| Fig. 4G: mean duration of social investigation bout, comparing bouts overlapping with periods of laser stimulation vs. bouts non-overlapping<br>• Two-way ANOVA with repeated measures on one factor (between-subjects factor = group, within-subjects factor = laser overlap) | POA-social-ChR2, overlapping with laser:<br>5.75 ± 2.37 (N = 8)<br><br>POA-social-ChR2, non-overlapping with laser:<br>2.07 ± 0.88 (N = 8)<br><br>POA-social-GFP, overlapping with laser:<br>3.95 ± 2.14 (N = 6)<br><br>POA-social-GFP, non-overlapping with laser:<br>2.72 ± 0.75 (N = 6)<br><br>N = 1 ChR2 female excluded that did not have any social investigation bouts overlapping with periods of laser stimulation | Main effect of group: F(1,12) = 0.60, P = 0.45<br>Main effect of laser: F(1,12) =19.97, **P < 0.003**<br>Interaction: F(1,12) = 4.96, **P = 0.046**<br><br>Within group post-hoc pairwise comparisons:<br>POA-social-ChR2 overlapping with laser vs. non-overlapping: t(12) = 5.11, **P = 0.001**<br>POA-social-GFP overlapping with laser vs. non-overlapping: t(12) = 1.48, P = 0.47<br><br>Across group post-hoc pairwise comparisons:<br>POA-social-ChR2 overlapping vs. POA-social-GFP overlapping: t(12) = 1.47, P = 0.49<br>POA-social-ChR2 overlapping vs. POA-social-GFP non-overlapping: t(12) = 3.47, **P = 0.02**<br>POA-social-GFP overlapping vs. POA-social-ChR2 non-overlapping: t(12) = 1.93, P = 0.27<br>POA-social-ChR2 non-overlapping vs. POA-social-GFP non-overlapping: t(12) = -1.44, P = 0.50 |
| Fig. 5B: effects of chemogenetic inhibition of POA neurons on resident-initiated investigation, females GH during TRAPing<br>• Paired t-test | Saline:<br>535.6 ± 203.9 (N = 5)<br><br>CNO:<br>516.0 ± 220.1 (N = 5) | T(4) = 0.15, p = 0.88 |
| Fig. 5D: effects of chemogenetic inhibition | Saline:<br>2130 ± 1552 (N = 5) | T(4)= 1.22, p = 0.29 |

| | | |
|---|---|---|
| of POA neurons on total USVs, females GH during TRAPing<br>• Paired t-test | CNO:<br>1654 ± 1048 (N = 5) | |
| Fig. 5F, left: proportion of mice with non-zero USV rates, pre-laser vs. laser, solo sessions, females GH during TRAPing<br>• McNemar's test for paired proportions | Pre-laser: 0 of 6<br><br>During laser: 1 of 6 | X^2(1) = 0, p = 1.00 |
| Fig. 5F, right: proportion of mice with non-zero USV rates, pre-laser vs. laser, social sessions, females GH during TRAPing<br>• McNemar's test for paired proportions | Pre-laser: 2 of 6<br><br>During laser: 3 of 6 | X^2(1) = 0, p = 1.00 |
| Fig. 5G: percentage of laser stimulations followed by social investigation<br>• T-test | POA-social-ChR2, female GH during TRAPing:<br>41.5 ± 31.2 (N = 9)<br><br>POA-social-GFP:<br>10.8 ± 5.0 (N = 6) | t(10) = 0.21, p = 0.84 |
| Fig. 5J: effects of chemogenetic inhibition of POA neurons on resident-initiated investigation, females tested as GH<br>• Paired t-test | Saline:<br>38.3 ± 23.8 (N = 7)<br><br>CNO:<br>34.7 ± 25.1 (N = 7) | t(6) = 0.99, p = 0.36 |
| Fig. 5L: effects of chemogenetic inhibition of POA neurons on total | Saline:<br>137.0 ± 127.1 (N = 7)<br><br>CNO: | t(6) = 2.47, **p = 0.048** |

| | | |
|---|---|---|
| USVs, females tested as GH<br>• Paired t-test | 15.3 ± 14.8 (N = 7) | |
| Fig. 5M: effects of chemogenetic inhibition of POA neurons on resident-initiated investigation, females tested as SH<br>• Paired t-test | Saline:<br>389.9 ± 87.7 (N = 7)<br><br>CNO:<br>78.9 ± 29.3 (N = 7) | t(6) = 8.52, **p < 0.001** |
| Fig. 5N: effects of chemogenetic inhibition of POA neuron on resident-initiated mounting time, females tested as SH<br>• McNemar's test for paired proportions | Saline: 6 of 7<br><br>CNO: 0 of 7 | X^2(1) = 4.167, **p = 0.041** |
| Fig. 5O: effects of chemogenetic inhibition of POA neurons on total USVs, females tested as SH<br>• Paired t-test | Saline:<br>2136 ± 565.5(N = 7)<br><br>CNO:<br>45.0 ± 11.0 (N = 7)<br>Saline:<br>2136 ± 565.5(N = 7)<br><br>CNO:<br>45.0 ± 11.0 (N = 7) | t(6) = 9.72, **p < 0.0001** |
| Fig. 6A: total resident-initiated social investigation<br>• Two-way ANOVA (factor 1 = housing; factor 2 = social context) | Male-female, group-housed resident:<br>291.0 ± 152.2 (N = 8)<br><br>Male-female, single-housed resident:<br>455.1 ± 95.8 (N = 8)<br><br>Male-male, group-housed resident:<br>214.1 ± 73.8 (N = 7) | Main effect of housing: F(1,26) = 6.55, **P = 0.02**<br>Main effect of social context: F(1,26) = 13.74, **P < 0.001**<br>Interaction: F(1,26) = 2.92, P = 0.10 |

| | | |
|---|---|---|
| | Male-male, single-housed resident:<br>246.8 ± 68.5 (N = 7) | |
| Fig. 6B: proportion of trials with resident-initiated mounting<br>• Z-test for independent proportions | Male-female, group-housed resident: 2 of 8<br><br>Male-female, single-housed resident: 6 of 8<br><br>Male-male, group-housed resident: 1 of 7<br><br>Male-male, single-housed resident: 0 of 7 | Male-female, group-housed resident vs. single-housed resident:<br>Z = -2, **P = 0.046**<br><br>Male-male, group-housed resident vs. Single-housed resident:<br>Z = 1.04, P = 0.30 |
| Fig. 6C: total USVs<br>• Two-way ANOVA (factor 1 = housing; factor 2 = social context); post-hoc Tukey's HSD tests | Male-female, group-housed resident:<br>545.3 ± 282.5 (N =8)<br><br>Male-female, single-housed resident: 2076.9 ± 462.2 (N = 8)<br><br>Male-male, group-housed resident: 8.4 ± 7.9 (N = 7)<br><br>Male-male, single-housed resident: 21.1 ± 16.4 (N = 7) | Main effect of housing: F(1,26) = 56.29, **P < 0.001**<br>Main effect of social context: F(1,26) = 158.64, **P < 0.001**<br>Interaction: F(1,26) = 54.45, **P < 0.001**<br><br>Post-hoc pairwise comparisons<br>• MF group vs. MF single: t(26) = -10.89, **P < 0.001**<br>• MF group vs. MM group: t(26) = 3.69, **P = 0.006**<br>• MF group vs. MM single: t(26) = 3.60, **P = 0.007**<br>• MF single vs. MM group: t(26) = 14.21, **P < 0.001**<br>• MF single vs. MM single: t(26) = 14.12, **P < 0.001**<br>• MM single vs. MM group: t(26) = -0.09, P = 1.00 |
| Fig. 6D: total Fos-positive POA neurons<br>• Two-way ANOVA (factor 1 = housing; factor 2 = social context); post- | Male-female, group-housed resident: 229.9 ± 68.3 (N = 8)<br><br>Male-female, single-housed resident: 328.2 ± 69.5 (N = 8) | Main effect of housing: F(1,26) = 4.97, **P = 0.04**<br>Main effect of social context: F(1,26) = 5.52, **P = 0.03**<br>Interaction: F(1,26) = 4.91, **P = 0.04**<br><br>Post-hoc pairwise comparisons<br>• MF group vs. MF single: t(26) = -3.25, **P = 0.02** |

| | | |
|---|---|---|
| hoc Tukey's HSD tests | Male-male, group-housed resident: 227.0 ± 53.8 (N = 7)<br><br>Male-male, single-housed resident: 227.3 ± 42.7 (N = 7) | • MF group vs. MM group: t(26) = 0.09, P = 1.00<br>• MF group vs. MM single: t(26) = 0.09, P = 1.00<br>• MF single vs. MM group: t(26) = 3.24, **P = 0.02**<br>• MF single vs. MM single: t(26) = 3.23, **P = 0.02**<br>• MM single vs. MM group: t(26) = -0.01, P = 1.00 |
| Fig. 6F: effects of chemogenetic inhibition of male POA-social neurons on resident-initiated investigation<br>• Two-way ANOVA (factor 1 = group; factor 2 = drug) | Male POA-social-hM4Di, saline:<br>476.6 ± 88.4 (N = 10)<br><br>Male POA-social-hM4Di, CNO:<br>536.4 ± 208.6 (N = 10)<br><br>Male POA-social-GFP, saline:<br>292.8 ± 109.7 (N = 10)<br><br>Male POA-social-GFP, CNO:<br>325.0 ± 118.6 (N = 10) | Main effect of group: F(1,18) = 13.76, **P = 0.002**<br>Main effect of drug: F(1,18) = 2.05, P = 0.17<br>Interaction: F(1,18) = 0.18, P = 0.67 |
| Fig. 6G: effects of chemogenetic inhibition of male POA-social neurons on proportion of trials with resident-initiated mounting<br>• McNemar's test for paired proportions | Male POA-social-hM4Di, saline: 6 of 10<br><br>Male POA-social-hM4Di, CNO: 0 of 10<br><br>Male POA-social-GFP, saline: 9 of 10<br><br>Male POA-social-GFP, CNO: 9 of 10 | Male POA-social-hM4Di, CNO vs. saline: X^2 (1) = 4 167, p = **0.04**<br><br>Male POA-social-GFP, CNO vs. saline: X^2 (1) = 0 167, p = 1.0 |
| Fig. 6H: effects of chemogenetic inhibition of male POA-social neurons on total USVs<br>• Two-way ANOVA (factor | Male POA-social-hM4Di, saline:<br>1868 ± 1097.4 (N = 10) | Main effect of group: F(1,18) = 0.35, P = 0.56<br>Main effect of drug: F(1,18) = 0.00, P = 0.96<br>Interaction: F(1,18) = 0.12, P = 0.74 |

| | | |
|---|---|---|
| 1 = group;<br>factor 2 = drug) | Male POA-social-hM4Di, CNO:<br>1826.4 ± 1120.5 (N = 10)<br><br>Male POA-social-GFP, saline:<br>1587.3 ± 764.1 (N = 10)<br><br>Male POA-social-GFP, CNO:<br>1643.9 ± 674.6 (N = 10) | |
| Fig. S5A: comparison of TRAPing session resident-initiated social investigation for POA-social-hM4Di groups<br>• T-test | Male POA-social-hM4Di:<br>365.8 ± 140.0 (N = 8)<br><br>Female POA-social-hM4Di:<br>553.9 ± 194.7 (N = 16) | t(22) = 2.43, **P = 0.02** |
| Fig. S5B: comparison of TRAPing session mounting for POA-social-hM4Di groups<br>• Mann Whitney U test | Male POA-social-hM4Di:<br>190.5 ± 154.9 (N = 8)<br><br>Female POA-social-hM4Di:<br>23.3 ± 45.6 (N = 16) | Z = -2.88, **P = 0.004** |
| Fig. S5C: comparison of TRAPing session USVs for POA-social-hM4Di groups<br>• T-test | Male POA-social-hM4Di:<br>1678 ± 1024 (N = 8)<br><br>Female POA-social-hM4Di:<br>2534 ± 637.1 (N = 17) | t(23) = -2.03, P = 0.054 |
